# Hormonal Interventions for optimizing reproductive efficiency in pigs at Farmers’ field

**DOI:** 10.1101/2023.02.06.527245

**Authors:** Sunil Kumar, Rafiqul Islam, Santanu Banik, Keshab Barman, P.J. Das, N.H. Mohan, Satish Kumar, Vivek Kumar Gupta

## Abstract

**Objective:** Optimizing reproductive efficiency in terms of estrus induction and production performance using simplified hormonal intervention using GnRH, gonadotropins and progesterone at farmers’ field conditions to bred animals with superior quality semen and also to reduce the maintenance cost.

**Method:** **T**he present study was planned for optimizing reproductive efficiency using four different easily available hormonal agents with simplified protocols at farmers’ field. A total of 126 females were used in the study. Hormonal protocols I, II, III and IV using GnRH + (PMSG+ hCG), (PMSG+ hCG), PMSG alone and prepared progesterone gel respectively were used for estrus induction in gilts and sows.

**Results:** The estrus induction/rate was (77.77%, 81.36%, 78.57% and 50% respectively. The corresponding figures for interval (hrs) of heat exhibition from hormonal administration (121.33±4.93, 121.54±3.60, 78.52±4.52 and 192±24), conception rate (%) (72.22%, 81.25%, 68.18% and 33.33%) farrowing rate (55.55%, 68.29%, 68.18% and 33.33 %), total litter size at birth (7.2±0.64, 7.90±0.47, 8.90±0.47 and 7.00±0) and hormonal cost (Rs.) per animal (570, 335, 280 and 250 Rs.) were respectively. The easier to use protocol was III followed by II, I and IV.

**Conclusion:** It was found that protocol II and III are effective and easier so can be used for optimizing reproductive efficiency in pigs at farmers’ field.

## INTRODUCTION

Reproductive failure is the most common reason for culling gilts and sows (Knauer *et al*., 2006; Tummaruk *et al*., 2009). Approximately 30% of replacement gilts never farrow (Stancic *et al*., 2011), because gilts fail to become cyclic or show delayed cyclicity (Saito *et al*., 2011). Initiation of puberty is dependent upon the activation of the hypothalamic– pituitary–gonadal axis. Puberty comes with activation of high-frequency pulses of LH (Diekman *et al*., 1983; Lutz *et al*., 1984; Camous *et al*., 1985) that are driven by cyclic increases in the secretion of gonadotropin-releasing hormone (GnRH) from the hypothalamus (Lutz *et al*., 1985; Kraeling and Barb, 1990). Inadequate gonadotropin secretion may result in pubertal failure and contributes to extended weaning to estrus intervals in postpartum sows (Edwards and Foxcroft, 1983). Moreover, insufficient gonadotropin secretion can be responsible for seasonal infertility in gilts and sows (Armstrong *et al*., 1986; Barb *et al*., 1991). To counter such problems, gonadotropins and progesterone used for estrus induction and synchronization in pigs which regulate the events leading to follicular maturation and ovulation or altering the luteal phase. PG600 (MSD Animal Health, USA) contains pregnant mare serum gonadotropin (PMGS) and human chorionic gonadotropin (hCG). PMSG and hCG mimic FSH and LH actions respectively. Puberty in gilts can be induced using PG600 (Martinat-Botté *et al*., 2011) to initiate estrus or heat in non-cycling pre-pubertal gilts that are nearing their natural initiation of puberty (Breen *et al*., 2006). PG600 reported to decrease the return to estrus interval in weaned sows (Breen *et al*., 2006).

GnRH and its analogues has also been used for stimulation and synchronization of ovulation in pigs (Webel and Rippel, 1975; Von Kaufmann and Holtz, 1982; Brüssow *et al*., 1990; Baer and Bilkei, 2004; Taibl *et al*., 2007). GnRH has been used for both induction and synchronization of ovulation in gilts and sows (Bruss□w *et al*., 1996; Martinat-Botté *et al*., 2010). Several protocols using GnRH agonists were proposed for swine (Von Kaufmann and Holtz, 1982; Brüssow *et al*., 1990; Knox *et al*., 2003). GnRH as OvuGel (triptorelin acetate 0.1 mg/ml) stimulates ovulation 96 hour post weaned sow when used as vaginal gel and results in ovulation approximately 22 hours later (Knox *et al*., 2018).

Synthetic progesterone altrenogest oral preparations such as Matrix (0.22% w/v) and Regumate (0.4% w/v) when fed for 14-18 consective days to gilts return to heat 5-7 days after last feeding (Davis *et al*., 1987; Martinat-Botte *et al*., 1990; Estienne, 2003).

Use of PG600 and Ovugel are not licensed for use in some countries including India, therefore, PMSG (Folligon, Intervet) and hCG (Chorulon, Intervet) were used in the present investigation to mimic the use of PG600. Further, PG600 is not effective to induce estrus in gilts that have already reached puberty (begun to cycle). The lack of response to treatment with P.G.600 is most often associated with treating gilts that have already reached puberty. Once the gilts reached puberty but due to shortage or lack of breedable male or A.I. service at farmers’ field that cycle leads to anestrum in the gilt. Therefore, GnRH administration prior to combination of PMSG+hCG is tried in the current study. Further, to mimic the effect of PG600, preparations of PMSG and hCG needs to be mixed in required ratio which makes the application difficult and requires skill. Moreover, once the vials containing lyophilized PMSG and hCG are opened, it needs to be used completely. Owing to the holding of smaller number of pigs by most of the Indian Farmers it may not be possible to use the complete content of chorulon (hCG) vials. Hence, the use of PMSG alone was also studied in the present study. Further, the Ovugel is not approved for use in gilts. Further, literature documented protocols requires understanding of the reproductive or specifically the ovarian dating of the female before application of hormonal agents for estrus induction or synchronization. Knowing ovarian dating is difficult to reach under backyard farmers’ field conditions. Hence, the present study was planned for optimizing reproductive efficiency in terms of estrus induction and production performance using simplified hormonal intervention using GnRH, gonadotropins and progesterone at farmers’ field conditions to bred animals with superior quality semen and also to reduce the maintenance cost.

## MATERIALS AND METHODS

### Animal Care

The study was conducted at the Indian Council of Agricultural Research-National Research Centre on Pig, Rani, Guwahati, 781131, India. The experimentation was carried out with prior approval from the Institute Animal Ethics Committee. The approved animal use protocol number is NRCP/CPCSEA/l658/IAEC-21 dated 23^rd^ July, 2018.

### Animals and experimental design

The study was conducted in tropical lower Brahmaputra Valley of Assam (India) where average rainfall is about 1700 mm per annum. The maximum temperature rises upto 39°C in July-August and minimum falls to 10°C in January. The study was conducted in nearby villages that are located within a radius of 2–80 km from ICAR-NRC on Pig, Rani (Assam). The present investigation was conducted in gilts (n=17) and 2^nd^ to 4^th^ parity sows (n=109). Animals were fed with locally available resources and concentrate feed as per availability. The gilts (>8 months; 8.67±0.16) and sows (>1 month of weaning; 1.81±0.10) were presented with the history of absence of estrus signs. The hormones used in the study such as Folligon (PMGS), Chorulon (hCG) and Receptal (GnRH analogue buserelin acetate) were purchased from Intervet India Pvt. Ltd., India). Commonly available progesterone (CDH Pvt. Ltd., New Delhi) was used to prepare a gel in methyl cellulose and alcohol for intra-vaginal use. Total four hormonal protocols (I,II,III and IV) were applied. Day of mineral supplementation and deworming (Fenbendazole @10mg/kg b.wt. P.O.) was considered as day 0 for each protocol.

- In protocol I (n=23; 17 gilts and 6 sows), each animal was injected (I/M) with GnRH (Buserelin acetate @10 µg/animal) on 7^th^ day and combined dose of PMSG (400 I.U.) and hCG (200 I.U.) on 9^th^ day.
- In protocol II (n=59 sows), each animal was injected (I/M) with combined dose of PMSG (400 I.U.) and hCG (200 I.U.) on 9^th^ day. This protocol was used as control protocol.
- In protocol III (n=28 sows), each animal was injected (I/M) with PMSG (400 I.U.) alone on 7^th^ day.
- In protocol IV (n=6 sows), progesterone (20mg/kg b.wt; progesterone dissolved in 1% suspension of methyl cellulose and alcohol) based gel was prepared in fresh and used as intra vaginal application on 7^th^ day. The gel was inserted with help A.I. Catheter (IMV India Pvt. Ltd.)

Subsequent to the day of injection, females were observed for expression of heat signs. Estrus expression was confirmed by visual signs such reddening and swelling of vulva along with positive back pressure test. Single time artificial insemination (A.I.) with extended semen was carried out after 24h of standing reflex. The conception was confirmed with transabdominal ultrasonography (miniScan, Meditech, USA) at 35-50 days post insemination (Fig.1). Parameters estimated for comparison of protocols were estrus induced animals (%), interval for onset of heat from time of hormonal application (hrs), conception (%), farrowing rate (%), total litter size and cost of protocol (Rs.). The easy to use score (lowest to highest; +, ++, +++, ++++) of protocols was estimated based on the mixing procedure of hormones, number of times of hormonal administration required and post administration complication. Farrowing rate was defined as the proportion of females served that farrow. Conception percentage was calculated as proportion of females detected conceived out of the successfully estrus induced and inseminated animals. Data was analyzed using standard statistical procedures and student’s t test.

**Fig 1.**
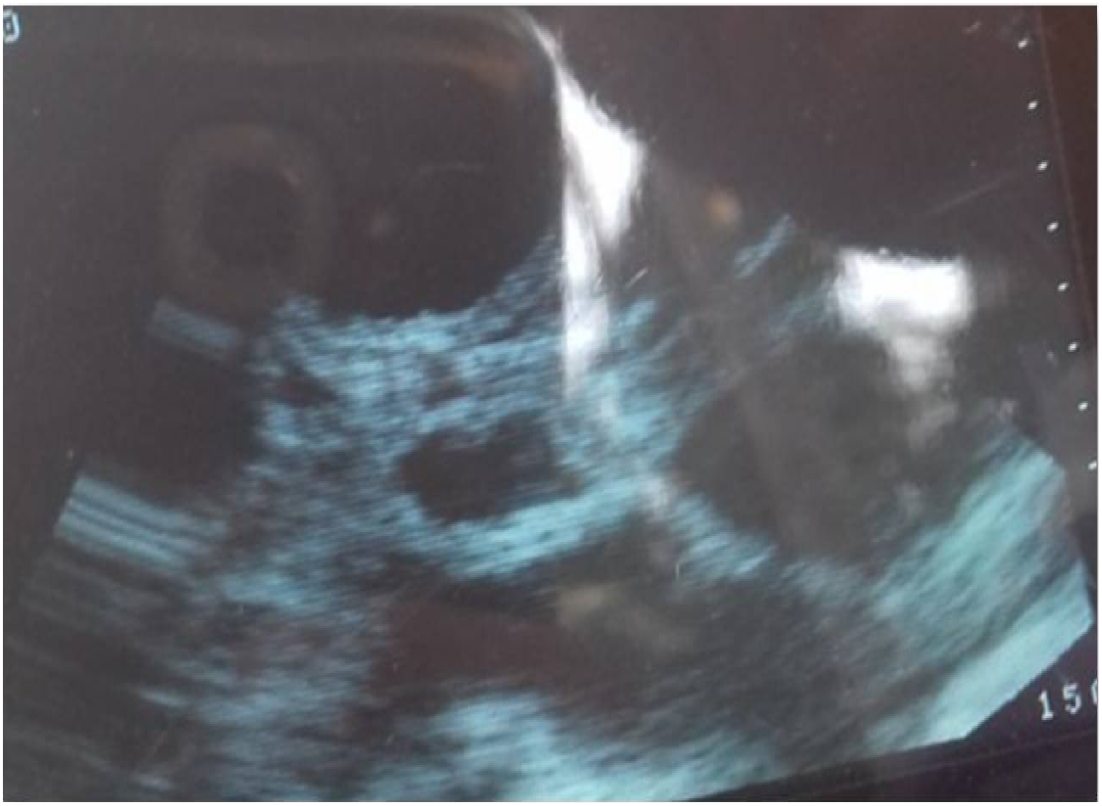
Representative image for pregnancy diagnosis using ultrasonography at 30-35 days in experimental animals. The black anechoic area surrounded by hyperchoic area are amniotic vesicles having conceptus inside them. It indicates that animal is pregnant.

## RESULTS

The estimation of different parameters in response to different hormonal treatment protocols is presented in Table 1. The overall estrus induction rate (%)in all protocols applied was 72.22% (91/126). Protocol II was the most successful (81.36%) for estrus induction followed by protocol III (78.57%), I (77.77 %) and IV (50%).The interval (h) from the administration of hormonal treatment to onset of estrus was 121.33±4.93, 121.54±3.60, 78.52±4.50 and 192±24 in protocols I, II, III and IV, respectively. Upon ultrasonographic pregnancy detection, the conception rate (%) in estrum served animals was highest in protocol II (81.25), followed by protocol I (72.22%), III (68.18%) and it was least in protocol IV (33.33%). The farrowing rate (%) was almost similar in protocol II (68.29%) and protocol III (68.18%). However, less farrowing rate was observed in protocol I (55.55%) and protocol IV (33.33%). The production performance in terms of total litter size at birth was, highest in protocol III (8.90±0.47) followed by II (7.90±0.47), I (7.2±0.64) and IV (7.00±0), respectively. The cost occurred for hormonal agents were higher in protocol I (570Rs.), followed by II (335Rs.), III (280Rs.) and IV (250Rs.). The ease of protocol in terms of field applicability, hormonal agents availability, preparation of dose was maximum in protocol III, followed by II, I and IV. Post treatment vaginitis was observed (2/6) in protocol IV animals. No anaphylactic shock or symptoms were observed after application of GnRH, PMSG and hCG in any protocol. Protocol III was observed as the simplest and cheaper to other three protocols (I, II, IV). In the protocol I, 82.35% gilts (14/17) and 66.66% sows (4/6) showed induced estrus response. The interval (h) from the hormonal treatment to exhibition of estrus in gilts and sows was (121.71±5.88) and (132±15.49), respectively. The conception (%) in gilts (9/14) and sows (4/4) was 64.28% and 100% respectively. However, the farrowing rate was very low 33.33 % in gilts (6/14) but in sows (4/4), the corresponding success was 100%. The litter size at birth in gilts and sows was (5.83±0.30) and (9.25±0.75), respectively.

**Table 1.** Estrus induction and production performance of animals with hormonal interventions at farmers’s field (mean ± SE)

| Parameters | Protocol I |  |  | Protocol II | Protocol III | Protocol IV |
| --- | --- | --- | --- | --- | --- | --- |
|  | Gilts | Sows | Total |  |  |  |
| No of animals treated | 17 | 6 | 23 | 59 | 28 | 6 |
| Estrus Induced (%) | 82.35<br>(14/17) | 66.66<br>(4/6) | 77.77<br>(18/23) | 81.36<br>(48/59) | 78.57<br>(22/28) | 50<br>(3/6) |
| Interval (h) from hormone administration to heat onset | 121.71±5.88 | 132±15.49 | 121.33±4.93 | 121.54±3.60 | 78.52±4.50 | 192±24 |
| Conception (%) | 64.28<br>(9/14) | 100<br>(4/4) | 72.22<br>(13/18) | 81.25<br>(41/48) | 68.18<br>(15/22) | 33.33<br>(1/3) |
| Farrowing rate (%) | 33.33<br>(6/14) | 100<br>(4/4) | 55.55<br>(10/18) | 68.29<br>(28/41) | 68.18<br>(15/22) | 33.33<br>(1/3) |
| Total Litter Size at birth | 5.83±0.30 | 9.25±0.75 | 7.2±0.64 | 7.90±0.47 | 8.90±0.47 | 7.00±0 |
| Hormonal Cost/animal(Rs.) | 570 | 570 | 570 | 335 | 280 | 250 |
| Easy to use | ++ | ++ | ++ | +++ | ++++ | + |

## DISCUSSION

Poor follicle development is a leading factor associated with anestrus or delayed estrus expression and poor response to ovulation induction. Further, the physiological and behavioral changes associated with the estrous cycle are controlled by hormones. Lack or deficiency of hormonal level may result lack of estrus behavior or ovulation. Lack of estrus in gilts may be primary (the animal may be genetically or physically incapable of producing eggs) or secondary, caused by physical or behavioural factors during rearing which suppress normal estrus behaviour. Sows that do not show signs of heat in the first 10 days after weaning called as anestrus sows, which leads to a lengthening of the wean-to-service interval, with consequent production and economic repercussions. Estrus Induction in such gilts and sows can be of great benefit to minimize the non productive days in swine production. However, the response and effectiveness, outcome and benefits of the estrus induction technique depend on ovarian dating, hormonal agent administered, regimen, management and production system (Estienne and Hartsock, 1998; Estill, 2000; Breen *et al*., 2005; Patterson *et al*., 2010). In the present study, the comparative efficacy of different simplified hormonal protocols for estrus induction and subsequent production performance was estimated at farmers’ field conditions.

GnRH used prior to combination of PMSG and hCG in protocol-I resulted a similar estrus response (77.77%) but lesser farrowing rate (55.55%) and litter size (7.2±0.64). In literature, it was found that treated sows received 10 μg buserelin after weaning and were inseminated once 30-33 hours later to get a farrowing rate of 87% and litter size of 13.6 ± 3.8 (Driancourt *et al*., 2013). Further, effect of 150 μg gonadrelin at breeding on reproductive performance of gilts resulted a farrowing rate of 88.9% with total litter size of 10.4 ±0.3. In protocol I, GnRH use prior to PMSG +hCG was based on the assumption that field animals might be having inactive ovaries due to poor mangemental conditions and negative energy balance. Subsequently, there might be unavailability of follicles for action of FSH and LH. Further, the mechanisms by which GnRH and hCG induces ovulation are different. GnRH induces ovulation by entailing a preovulatory peak of LH, while hCG acts directly on LH receptors located on ovarian follicle cells (Wongkaweewit *et al*., 2012). Therefore, it is possible that, under stress (lactational, negative energy balance, summer etc.), some sows might have inadequate pituitary LH stored to mount an effective ovulatory surge in response to GnRH treatment. The GnRH used in protocol-I might have caused the follicle development and growth of small follicles to medium sized follicles in such anestrum animals. Then, such medium follicles might have developed LH receptors for action of exogenously administered hCG on 9^th^ day. Further, there is species variation in the interplay actions of estrogen, Kisspeptin and GnRH to triger LH surge for estrus (Kraeling and Barb, 1990; Britt *et al*., 1991; Estienne *et al*., 1997; Casper, 2015). In comparison to other livestock species, much success has not been reported for improving reproductive efficiency using GnRH in Pig. This may be due to anatomical and physiological differences in pig as swine are the only livestock species that produce both the second mammalian isoform of gonadotropin-releasing hormone (GNRH2) and its receptor (GNRHR2). Indeed, GNRH2 is a poor stimulator of gonadotropin release; GNRH2-induced secretion of luteinizing hormone (LH) and follicle-stimulating hormone (FSH) is about 10% that of GNRH1 (Millar *et al*. 1986; Desaulniers *et al*., 2017). Further, it was found that the number of KISS1-expressing cells in the PeN but not in the ARC was significantly upregulated by estradiol treatment in ovariectomy pigs (Tomikawa *et al*., 2010a) suggesting that the anterior population of kisspeptin neurons is involved in LH surge generation in pigs as well. Moreover, kisspeptin neurons are mainly located in the hypothalamic periventricular nucleus in pigs (Tomikawa *et al*., 2010b) while in ruminants it is in preoptic area of hypothalamus. In the last decade, kisspeptin has emerged as being critically important in controlling gonadotropin secretion and is central to the effects of nutrition, disease, stress and season on gonadotropin secretion in laboratory and livestock species. Further research work in the area of GnRH and kisspeptin application is needed in pig reproductive biology which will provide further opportunity for augmenting reproductive efficiency in pigs.

The estrus response observed using protocol II (81.36%) and III (78.57%) was found lesser than in a previous (86.4%) report (Kadirvel *et al*., 2017) under Indian conditions. Kadirvel et al. (2017) found that 86.4% of sows in the treatment group exhibited estrus in response to administration of PMSG (800IU) and hCG (500IU) where the average interval between treatment and onset of estrus was 84.8±2.43 hours in comparison to 121.54±3.60 hrs in protocol II and 78.52±4.50 hrs in protocol III. Subsequently, in the previous study, farrowing rate of 82.6% and litter size of 9.2 ± 0.32 after estrus synchronization with timed insemination was reported in comparison to protocol II (68.29; 8.90±0.47) and protocol III (68.18; 7.90±0.47). The lower farrowing rate and production performance in the current study may be due to different managental conditions during pregnancy and also might be due to lesser quantity of hormones used. In a foreign study using PG 600, Breen et al. (2006) reported that estrus response was 98.2 %, along with acceptable conception rate (92.9%) and farrowing rate (76.4 %). Further, Kirkwood et al. (1998) evaluated performance of gilts bred at a PG600®-induced estrus response in 78%, time taken in onset of estrus 39.8 ±1.4 hrs, farrowing rate (74.4%) and total Litter size (9.4 ±0.3). PMSG and hCG successfully induced estrus in experimental animals. The basis of success is supported by the fact that that FSH is necessary to support follicle development beyond 2-3 mm and LH beyond 4 mm (Driancourt *et al*., 1995). Once the follicles reach a larger preovulatory stage FSH declines (Guthrie and Bolt, 1990) and LH pulse secretion changes from luteal (low frequency) to follicular patterns (high frequency). Final stage of follicular development is associated with decreased FSH and increased LH receptor expression along with increased production of estradiol, as well as increased fluid accumulation within the follicular cavity. Increased preovulatory estradiol finally elicits the GnRH-mediated LH release from the pituitary, thereby initiating ovulation and the release of a mature oocyte. The PMSG and hCG treatment in the experimental might have given a boost to existing smaller, quiescent stage to functional and ovulatory follicles. It may be confirmed from the present study that the ready to use commercially available PG600 and prepared combination of PMSG and hCG has more or less similar effect in terms of farrowing rate.

In the present study, the use of PMSG alone (Protocol III) was found effective for estrus induction (78.57%), that may be due to possible follicular maturation. The finding is supported by previous reports where single injections of eCG with or without hCG induced follicle maturation and estrus expression within a 4 to 5 day period (Hühn *et al*., 1996; Brüssow *et al*., 1996; Brüssow *et al*., 2009; Bennett *et al*., 2008; Driancourt, 2013). From a practical standpoint, eCG alone and eCG and hCG combinations require a single injection, and can induce estrus in 60%–70% of prepubertal gilts and improve estrus expression in weaned sows by 10%–15%, depending upon parity, lactation length, and season (Knox, 2015).

The successful use of PMSG alone may be due the fact that there might be the possibility of the medium sized follicles in the experimental animals but such follicles were unable to grow further due to lack of sufficient gonadotropin stimulation. The facts are supported by the reports where it was concluded that small follicles bind predominantly FSH and show maximal FSH receptor gene expression while medium-sized follicles can bind both FSH and LH, and expresses the genes for each type of receptor (Knox, 2018). Further, in large follicles, LH binding and receptor gene expression is high, while FSH binding is minimal and FSH receptor gene expression is not detectable (Lucy *et al*., 2001; Ainsworth *et al*., 1990).

In the protocol IV, prepared progesterone gel for intravaginal use was not found much effective. Intention was to identify the similar progesterone effect as used in bovines (eg. CIDR, PRID etc.) for estrus induction and synchronization. This is because, currently progesterone analogue preparation (Matrix and Regumate) and GnRH based Ovugel are not available worldwide. Available oral progestagens feeding based preparations have limitations of daily feeding and accurate dose determination at farmers’field.

The differential fertility responses in used protocols (I,II,III and IV) might have been affected by lactation length, body condition score, back fat thickness, season, parity and required adjustments in the timing of hormone administration used for follicle induction, maturation and ovulation. The success of exogenous ovulation induction in pigs appears to depend upon the hormonal agent, dose, stage of follicle development at time of treatment, and ensuring that treatment occurs prior to the endogenous LH surge. In the animals where estrus induction response was not successful, it may be due to failure to induce follicle development and subsequent lack of estrus expression may be associated with the ratio of FSH:LH activity, the concentration of the hormones, and the duration of stimulation used in the protocols. Further, it may be due to the fact that hormonal administration too early in the stage of follicle development may ovulate more immature follicles, induce cysts, and inhibit estrus expression (Nissen *et al*., 1995). There may be other reasons in females failed to respond to applied protocols, which may include problems of inactive ovaries, cystic follicles, presence of corpora lutea with or without inapparent genitourinary disease and reproductive tract abnormalities.

Lack or restriction of follicular makes the failure of estrus exhibition or ovarian cyclicity. Therefore, FSH, LH and GnRH have important role in follicular growth, ovulation and estrus exhibition. These hormones are used across the world except some countries where such hormonal agents are not licensed for sale and use. Further, it is critical to clearly understand the estrous cycle of pigs before applying specific method for estrus induction or synchronization. However, at farmers’ field and more particularly backyard pig farming, it makes difficult to apply estrus induction or synchronization protocols. The use of male effect by introduction of mature boar in a group of gilts provides opportunity for bringing gilts in estrus. However, this practice is usually undertaken in organized farms in intensive farming. Introduction of male is not easy in unorganized backyard system where there is lack of mature boar availability. This is because, keeping and maintaining a mature boar is a costly affair for farmer under tropical conditions. Hence, farmers become reluctant to keep boars, subsequently it results indirect loss in terms of delayed puberty or anestrum in gilts. Similar condition of anestrum exists in case of backyard sows where piglets are usually kept with nursing mother for longer duration than recommended period or piglets are not weaned till the sale of piglets to get income. Ultimately it results a prolonged lactational anestrum in sows where body condition of mother also become a questionable event. During lactation and before weaning, follicles exhibit a wave-like pattern of growth, and emerge from a cohort of 20-30 follicles of 2 mm that grow to not larger than 5 mm (Lucy *et al*., 2001). This is because suckling inhibits secretion of GnRH and subsequently LH due to a concerted action of prolactin, oxytocin and endogenous opioid peptides that prevent final growth of these follicles to reach ovulatory size before weaning (Varley and Foxcroft, 1990). Once the piglets are weaned, follicles start to grow to 7-8 mm before ovulation. Initially FSH increases and then decreases at weaning. Basal LH and LH pulse frequency increases; these are key regulators of postweaning follicle development which affect the weaning to estrus interval (van de Brand *et al*., 2000). Hence, if hormonal agents are applied with proper management, such economic losses can be minimized. However, it is difficult to understand the ovarian or estrus cycle status of female under farmers’ field condition in backyard system because of lack of awareness, poor management, and lack of resources. Additionally, it becomes difficult to choose the type of hormonal protocols to be applied for the presented female. Hence, combination of PMSG+hCG (Protocol II) and PMSG alone (Protocol III) for the estrus induction in gilts and sows with unknown ovarian status may be used at farmers’ field.

## CONCLUSION

In conclusion, Protocol II (PMSG @ 400 I.U. and hCG @ 200 I.U. per animal) and Protocol III (PMSG alone @ 400 I.U. per animal) can be used for estrus induction and optimizing subsequent production performance effectively under farmers’ field condition. The PMSG alone use is easier and cost effective under pig backyard farming. Use of GnRH prior to gonadotropins administration and intravaginal progesterone found not much useful for application. Although research efforts were made in past for developing estrus induction and synchronization protocols in pigs, yet much work is needed from practical application view at farmers’ field. The present study provides a good opportunity to explore further basis of estrus induction and synchronization for the purpose of optimizing reproductive efficiency in pigs.

## ACKNOWLEDGMENTS

The authors are grateful to the Director, ICAR-National Research Centre on Pig, Rani, Guwahati, 781131, India for providing the facility for conduction of the experiment.

